# Knock out of the intracellular calcium conducting ion channel Mitsugumin 23 (MG23) protects against pressure overload induced left ventricular hypertrophy and cardiac dysfunction

**DOI:** 10.1101/2024.06.28.601299

**Authors:** Amy M. Dorward, Gavin B. Robertson, Claire Sneddon, Chloe L. O’Rourke, In Hwa Um, David J. Harrison, Miyuki Nishi, Hiroshi Takeshima, Colin E. Murdoch, Samantha J. Pitt

**Author notes:** Corresponding author(s): Dr Samantha J. Pitt, School of Medicine, Medical and Biological Sciences Building, University of St Andrews, St Andrews, Fife, KY16 9TF, United Kingdom. Dr Colin E. Murdoch, Division of Cellular and Systems Medicine, School of Medicine, Jacqui Wood Cancer Centre, James Arrott Drive, Ninewells Hospital & Medical School, Dundee, DD1 9SY, United Kingdom. Both authors contributed equally to this work (Amy M. Dorward, Gavin B. Robertson).

## Abstract

**Background:** In cardiac dysfunction, intracellular Ca^2+^-dynamics are disrupted leading to leakage of Ca^2+^ from the sarcoplasmic reticulum (SR). This results in diminished cardiac contractility and impaired cardiac function. In cardiac tissue, the underlying molecular mechanisms responsible for RyR2-independent Ca^2+^ leak are poorly understood. Mitsugumin 23 (MG23) is an intracellular Ca^2+^-conducting ion channel located on ER/SR and nuclear membranes. We propose that MG23 contributes to regulation of intracellular Ca^2+^-homeostasis, and that altered MG23 function may drive progression of cardiac dysfunction. The aim of this research was to investigate the role of MG23 in SR Ca^2+^ leak, and whether knock out of *Mg23* protects the heart against pressure-overload induced left ventricular hypertrophy.

**Methods:** Cardiac pressure-overload was induced in wild type (WT) and *Mg23*-knock out (KO) mice through subcutaneous Angiotensin II (AngII, 1.1 mg/kg/day) infusion via osmotic pump. After 10-days infusion, *in vivo* pressure-volume dynamics were measured by insertion of a pressure-volume catheter into the left ventricle. MG23 protein expression was assessed through Western blot analysis. Ventricular fibrosis and cardiomyocyte size were measured using histological and immunofluorescence approaches. Cardiomyocytes were isolated from WT and *Mg23*-KO hearts and intracellular Ca^2+^ dynamics assessed through live cell imaging using the Ca^2+^ indicator Fluo-4.

**Results:** AngII-induced cardiac pressure-overload increased expression of MG23 in WT mouse hearts. Knock out of *Mg23* protected hearts against AngII-induced cardiac hypertrophy. Compared to WT animals, AngII treated *Mg23*-KO mice displayed a significant reduction in left ventricular fibrosis and displayed normal cardiac functioning. In *Mg23-*KO hearts, no alteration in expression of key Ca^2+^ handling proteins was identified, but cardiomyocytes displayed altered Ca^2+^ spark profiles consistent with a role for MG23 in SR Ca^2+^ leak.

**Conclusion:** MG23 plays a key role in driving Ca^2+^ dysregulation observed in the early pathological stages of pressure-overload induced heart failure.

## Introduction

Cardiac hypertrophy is an adaptive physiological mechanism that develops in response to hypertension associated with increased pressure overload of the heart and subsequent left ventricle remodelling to maintain cardiac function. If left untreated, cardiac hypertrophy can deteriorate into heart failure with elevation of cardiac fibrosis and dysregulation of calcium ion (Ca^2+^) handling. Several major signalling pathways involved in the cardiac hypertrophic response are activated by an increase in intracellular Ca^2+^ concentration^1^ but the origin(s) of this subcellular Ca^2+^ signal are not fully understood.

Cardiac contractility is dependent on the controlled synchronous release of Ca^2+^ from sarcoplasmic reticulum (SR) intracellular stores mediated through a process called excitation-contraction (EC) coupling. Ca^2+^ influx into the cardiomyocyte is sufficient to activate clusters of the type-2 ryanodine receptor (RyR2) in discrete Ca^2+^-release events termed Ca^2+^-sparks^2^. Macroscopic summation of Ca^2+^-sparks leads to a transient rise in cytosolic [Ca^2+^] and initiation of cardiac contraction^3,4^.

Efficient termination of elevated intracellular Ca^2+^-concentrations is achieved through a balanced combination of Ca^2+^-extrusion from the cell through the Na^+^/Ca^2+^ exchanger and reuptake of Ca^2+^ into the SR through the SR/ER Ca^2+^-ATPase (SERCA) (for review see^5^. Although most RyR2-channels are closed during diastole, a small but important leakage of Ca^2+^ from SR-stores occurs^6^. This basal leak plays an important physiological role in subtle sensitisation of RyR2 for activation through Ca^2+^ induced Ca^2+^ release^7^ and protects the heart against spontaneous Ca^2+^-release caused by SR-store Ca^2+^-overload^8^. However, SR Ca^2+^-leak can become pathogenic. In heart failure, exacerbated diastolic SR Ca^2+^-leak is attributed to progressive deterioration of cardiac function^9–11^, implying dysfunction in mechanisms controlling Ca^2+^-flux across the SR membrane^12–14^. Defective RyR2-channel function has been shown to increase SR Ca^2+^-leak^9^, but its relevance as a causative factor rather than phenotypic consequence is questioned^15^, with increasingly prominent roles for RyR2-independent leak demonstrated^10,16^.

In cardiac hypertrophy, the question remains controversial whether altered patterns of Ca^2+^ release and reuptake associated with excitation-contraction coupling affect hypertrophic signalling pathways and drive pathology. Work by Van Oort and co-workers^17^ have shown that pathological leak of Ca^2+^ from the SR through mutant RyR2 accelerates the development of cardiac hypertrophy and heart failure after pressure overload. Non-clustered or orphaned RyR2s and more recently IP3R over-expression^16^ are thought to play a key role in pathophysiological cellular restructuring in heart failure^18,19^. However, this does not account for any contribution through the RyR2 independent Ca^2+^ leak pathway.

Mitsugumin 23 (MG23) is a Ca^2+^-conducting non-selective cation channel located with abundance on ER/SR and nuclear membranes encoded by *TMEM109*^20,21^. Recent work has shown that MG23 plays a role in SR Ca^2+^ release in skeletal muscle^22^ but the role in cardiac muscle is largely unknown. We propose that MG23-mediated Ca^2+^-release becomes more apparent under pathophysiological conditions^10^ raising the question of whether MG23 plays a key role in the pathogenesis of cardiac hypertrophy and heart failure through increased SR Ca^2+^ leakage.

The aim of this study was to determine the role of MG23 in the development of pathological cardiac hypertrophy through diastolic SR Ca^2+^ leak. To study the specific effects of increased MG23-mediated Ca^2+^ release leading to early-stage heart failure, we used a MG23 knock out mouse and investigated dynamic cardiac functioning, cardiac fibrosis and hypertrophy using angiotensin II mediated pressure overload as our model.

## Methods

### Detailed methods are provided in the supplementary material. Reagents

Unless otherwise stated all chemicals were AnalaR or best available equivalent grade from Sigma-Aldrich (Dorset, UK). All solutions used were made in de-ionised ultrapure water (18.2 MΩ, TOC ≥ 5 ppb, MilliQ; Millipore, Hertfordshire, UK) and those for use in isolation of mouse cardiomyocytes and planar lipid bilayer experiments were filtered through a MCE Millipore membrane filter (0.45 µm pore). The pH of all solutions was measured using a Hannah pH 211 microprocessor pH meter (Hannah Instruments Ltd., UK).

### Ethical consideration

The institutional ethical committees of the University of St Andrews, University of Dundee and Kyoto University granted a favourable ethical opinion (SEC17021) for the use of experimental animals in this study. The care and sacrifice of animals used conformed to Directive 2010/63/EU of the European Parliament on the protection of animals used for scientific purposes as well as the United Kingdom Animals (Scientific Procedures) Act 1986. Animal work was carried out under project license P82006EDF (S.J. Pitt) and P92B6930F (C.E. Murdoch).

### *In-vivo* animal studies

*Mg23* knockout (*Mg23-*KO) mice were generated as described by^22^ on a pure B57BL/6J background. Genotyping was undertaken by PCR.

Male *Mg23-*KO mice were compared with WT littermates at 12 weeks of age. Osmotic mini-pumps (Model 1002, Alzet, Cupertino, CA) filled with either saline or Angiotensin II (AngII) were subcutaneously implanted and set to infuse Ang II at a rate of 1.1 mg/k/day for 10 days^23^. Left ventricular pressure–volume (PV) relations were measured with a 1.2F pressure-volume catheter and ADVantage system (Transonic, ON, Canada) inserted retrogradely into the left ventricle via the right carotid artery under isoflorane (1.5-2%) anaesthesia as previously reported^23^. Animals were immediately euthanised with hearts perfused with high potassium solution (5 mM) in PBS to arrest heart in diastole, for subsequent tissue collection.

### Histochemistry/Immunofluorescence

Tissue was fixed in 10% buffered formalin, processed to paraffin blocks, sectioned at 4 µm and mounted on charged microscope slides. Picro-Sirius Red (PSR) staining was used to histochemically stain fibrillar collagen^24^. To identify cardiac capillary density, Isolectin-B4 (1:200, I21414) immunofluorescence labelling was carried out followed by Alexa Fluor 488 secondary antibody (1:200, S11223). Wheat germ agglutinin (WGA) (1:150, RL-1022) staining was used to label the plasma membrane. Full methods can be found in supplementary material.

Sections were scanned using Zeiss Axio Scan Z1 slide scanner (Zeiss, Germany) at x20 objective magnification. The images were then analysed using HALO® image analysis platform (version.3.5.3577.140, Indica Labs) and HALO AI (version 3.5.3577). Isolectin-B4 positive vessels were segmented using AI drive DenseNetA1 V2 classifier, while cardiomyocyte size was analysed by calculating cell area.

### IF protocol – NFATs

Tissue was probed using a multiplex immunofluorescence protocol, carried out by incubation with primary antibody, followed by sequential incubation of rabbit secondary HRP antibody (Leica) and TSA incubation (NFATc2 1:200, PA596227/Cy5 TSA; Troponin T 1:1,500, BS-10648R/Cy3 TSA; NFATc1 1:3,000, PA590432/FITC TSA). The final antibody incubation (NFATc3 1:2,000, PA599546) was followed by Rabbit biotin (1:200, 656140) secondary and Alexa Flour 750 (1:100, S21384). To avoid cross contamination between antibodies, heat-induced stripping method was performed with pH 6 sodium citrate buffer between each primary antibody visualisation. Tissue was mounted with ProLong Glass Antifade Mountant with NucBlue (P36985) and scanned as before. Following image acquisition using Zeiss Axio scan z1, coverslips were removed, and WGA staining and scanning was carried out on the slides as detailed above. Both multiplexed immunofluorescent images and WGA images were registered and fused on Halo and analysed utilising the HALO AI highplex FL v.4.1.3 analysis algorithm. WGA and NucBlue channels were used to segment cytoplasmic and nuclear compartment (Fig S2), and FITC, CY3, CY5, and Alexa Fluor 750 channels, visualised NFATc1, Troponin, NFATc2, and NFATc3, respectively - were measured their intensity in both cytoplasmic and nuclear compartment using HALO AI custom nuclear segmentation plugin.

### Cardiomyocyte isolation

Adult mouse cardiomyocytes were isolated using an adapted Langendorff-free protocol as previously described^25^. For detailed methods see supplementary material. Tyrode’s solution used was (in mM): 5 KCl, 135 NaCl, 0.33 NaH_2_PO_4_, 10 HEPES, 5 glucose, 5 Na-pyruvate and 1 MgCl_2_ titrated to pH 7.4 with NaOH (from^26^); Collagenase buffer: 0.5 mg/ml collagenase (type I; C0103), 1.67 mg/ml bovine serum albumin (BSA) and 0.6 mg/ml protease (type XIV; P5147) diluted in Tyrode’s.

Isolated cardiomyocytes were reintroduced to Ca^2+^ by three rounds of 20 minutes sequential gravity settling in perfusion buffer containing 500 µM, and 1 mM CaCl_2_, respectively. Cells were stored in Tyrode’s solution containing 1 mM Ca^2+^ ^26^ and used within 2 hours of isolation.

### Live-cell imaging

Fluorescence microscopy experiments were conducted on live cardiomyocytes using an Olympus FV1000-IX81 confocal laser scanning microscope system controlled by Olympus Fluoview software (V4.0). Freshly isolated cardiomyocytes were loaded with 5 μM Fluo-4 AM (λex= 494 nm, λem= 506 nm; F14201, Thermo-Fisher, UK) for 20 minutes at room temperature. Cells were washed in indicator-free Tyrode’s solution and kept at room temperature until use. Whole-cell (xy) images and line-scans (xt) were viewed using 40x (NA 0.85) or 100x (NA 1.4; oil-immersion) objective lenses, respectively. Laser power settings (1% power; 488 nm excitation, 515 nm emission; argon-ion) were kept constant.

### SR Ca^2+^-load

Isolated cardiomyocytes from both WT and *Mg23*-KO mice were loaded with Fluo-4 AM in nominally Ca^2+^-free Tyrode’s solution. To assess SR Ca^2+^-load in quiescent cells, 10 mM caffeine was added to the bath and cellular Ca^2+^-responses recorded, with whole-cell images obtained at a frequency of 2 Hz. Image analysis was performed offline using ImageJ software (NIH, USA).

### Ca^2+^-sparks

Prior to recording spontaneous Ca^2+^-sparks, Fluo-4 AM loaded cardiomyocytes were superfused with Tyrode’s solution containing 1 mM Ca^2+^ and field-stimulated (30 seconds; 1 Hz) to achieve steady-state conditions. The line-scan mode (xt) was used to observe spontaneous Ca^2+^-sparks along the longitudinal axis of each cell at a frequency of 511 Hz. Spontaneous Ca^2+^ sparks were detected and analysed using the SparkMaster plugin^27^ developed for Image J software, in accordance with established spark criteria^28^.

### Tissue lysis

Left ventricular tissue taken from *in vivo* animal studies was lysed in cell lysis buffer (#9803S, Cell signalling) supplemented with protease inhibitor tablet (11697498001, Roche). Tissue was homogenised in a FisherBrand Bead Mill 24 with ceramic beads (19-645-3, Omni International). Heart tissue from *Mg23-*KO animals and WT littermates was homogenised in homogenisation buffer composed of 20 mM Tris-HCl (pH 7.4), 0.3 M sucrose and protein inhibitors, 1 mM PMSF, 1 µg/ml antipain, 1 µg/ml leupeptin, 1 µg/ml pepstatin A. Samples were centrifuged at 8,000 xg for 10 minutes and supernatant collected.

### Western blot

Left ventricular homogenates from *in vivo* studies were size-fractionated by SDS-PAGE on 4-12% Bis-Tris precast gels as previously described^10^. Blots were probed with MG23 (1:5,000, HPA011785) and secondary horseradish peroxidase-linked goat anti-rabbit (1:10,000, ab97051). Blots were visualised using a Fujifilm LAS-3000 detection system. Densitometry was performed using ImageJ software (NIH, USA), and was normalised to total protein (No-Stain™ Protein Labeling Reagent, A44449). Heart homogenates were size-fractionated by SDS-PAGE. Blots were probed with the following: MG23 (1:2,000, ab121349), RyR2 (1:5,000, a gift from Dr Masahiko Watanabe, Hokkaido University^29^), NCX1 (1:3,000, a gift from Dr Takahiro Iwamoto, Fukuoka University^30^), SERCA2 (1:2,000, sc-8095), JP2 (1:2,000, custom,^31^), Calsequestrin (1:2,000, PA1-913), GAPDH (1:5,000, G9545) and Cav1.2 (1:200, ACC-003). Membranes were developed using autoradiography film. Densitometry was performed using ImageJ software (NIH, USA).

### Statistical Analyses

Statistical analysis was performed using GraphPad Prism software (GraphPad; CA, USA). Analysis specified in figure legends. Unless otherwise stated, data is shown as individual data points with a bar for mean.

## Results

### *Mg23*-KO hearts are protected against hypertrophy following AngII-induced pressure-overload

To investigate if MG23 plays a role in the pathogenesis of cardiac hypertrophy, Angiotensin II (1.1mg/kg/day) osmotic pumps were implanted in WT and *Mg23*-KO mice to induce cardiac pressure-overload. Following 10-day treatment with AngII there was a significant increase in the expression of MG23 compared to saline control animals (Fig 1). We next assessed cardiac morphology in WT and *Mg23*-KO mice following AngII treatment (Fig 2). WT mice displayed enlarged hearts shown by a significant increase in heart weight/femur length. This was a result of increased left ventricular mass as shown by a significant increase in left ventricular weight/femur length. Whole heart weight increased from 7.5 ± 0.74 mg/mm to 9.51 ± 1.44 mg/mm (Fig 2A, p = 0.0032) and left ventricular weight increased from 5.43 ± 0.77 mg/mm to 7.33 ± 0.84 mg/mm (Fig 2B, p = 0.0008). In *Mg23*-KO mice, there was no significant change in whole heart weight or left ventricular mass following AngII treatment. Heart weight/femur length was 7.72 ± 0.75 mg/mm in *Mg23*-KO saline controls and 8.42 ± 0.66 mg/mm following AngII treatment (p = 0.929) and left ventricular weight/femur length was 5.81 ± 0.58 mg/mm for saline control and 6.76 ± 0.68 mg/mm in AngII treated *Mg23*-KO animals (p = 0.468). There was no significant change in right ventricular mass (RV/femur, Fig 2C), lung, liver or kidney weight (Figs 2D-2G) in WT or *Mg23*-KO animals after Ang II treatment compared to saline controls (see Table S1 for all organ weights). To investigate if cardiac hypertrophy in response to Ang II treatment in WT mice resulted in enlarged cardiomyocytes, left ventricular tissue slices were stained with WGA-Rhodamine which binds to cell membranes (Fig 2H) enabling cardiomyocyte size to be measured. No significant change in average cardiomyocyte (CM) size (Fig 2I) or cardiomyocyte density (Fig 2J) between any of the treatment groups was observed.

**Figure 1:**
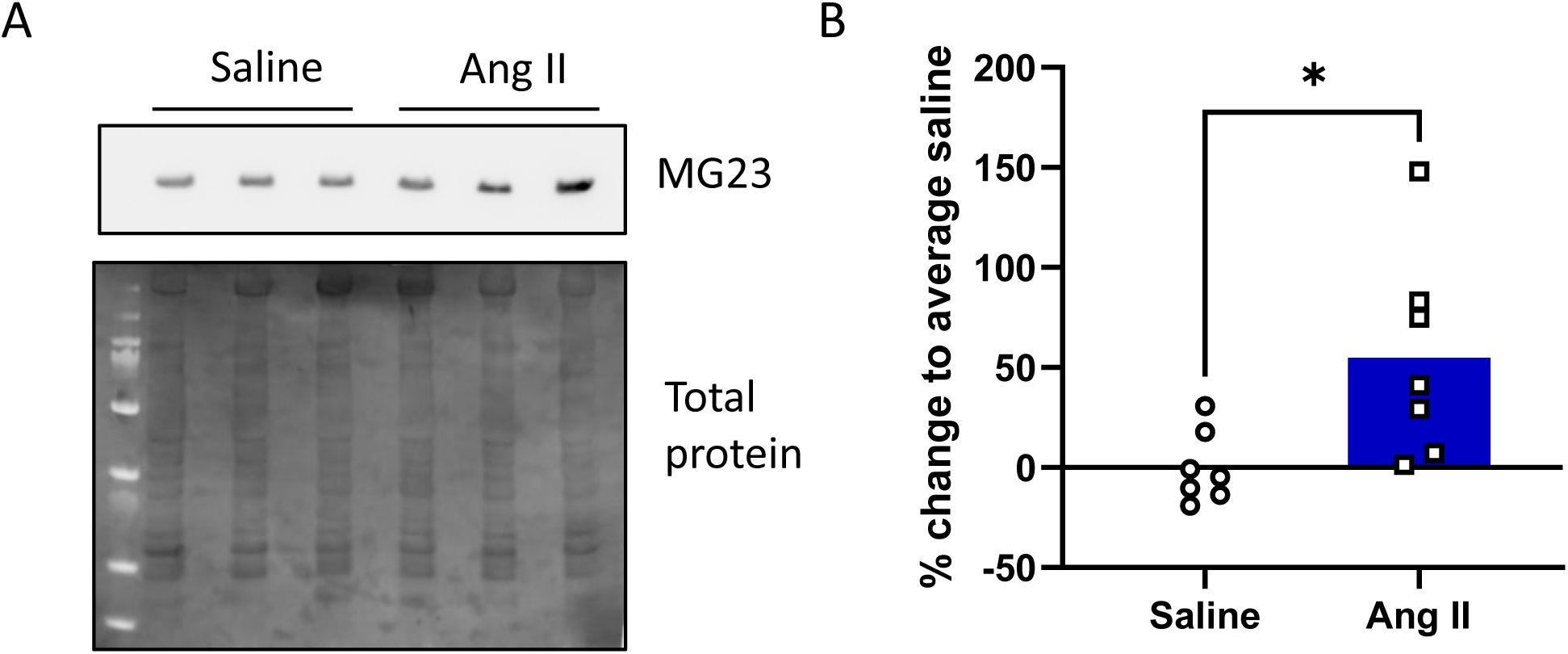
MG23 protein expression is increased in AngII-treated hearts. **A)** Representative Western blot of left ventricle heart homogenates. 30 µg protein per well. The upper panel shows MG23 protein expression while the lower panel shows total protein. **B)** Percentage change of MG23 expression in saline (black) and AngII (blue) relative to average saline control value. Data is displayed as individual points showing average expression per animal with a bar representing group average. n = 4 technical replicates. n = 7 animals per treatment. *p<0.05, unpaired t-test with Welch’s correction.

**Figure 2:**
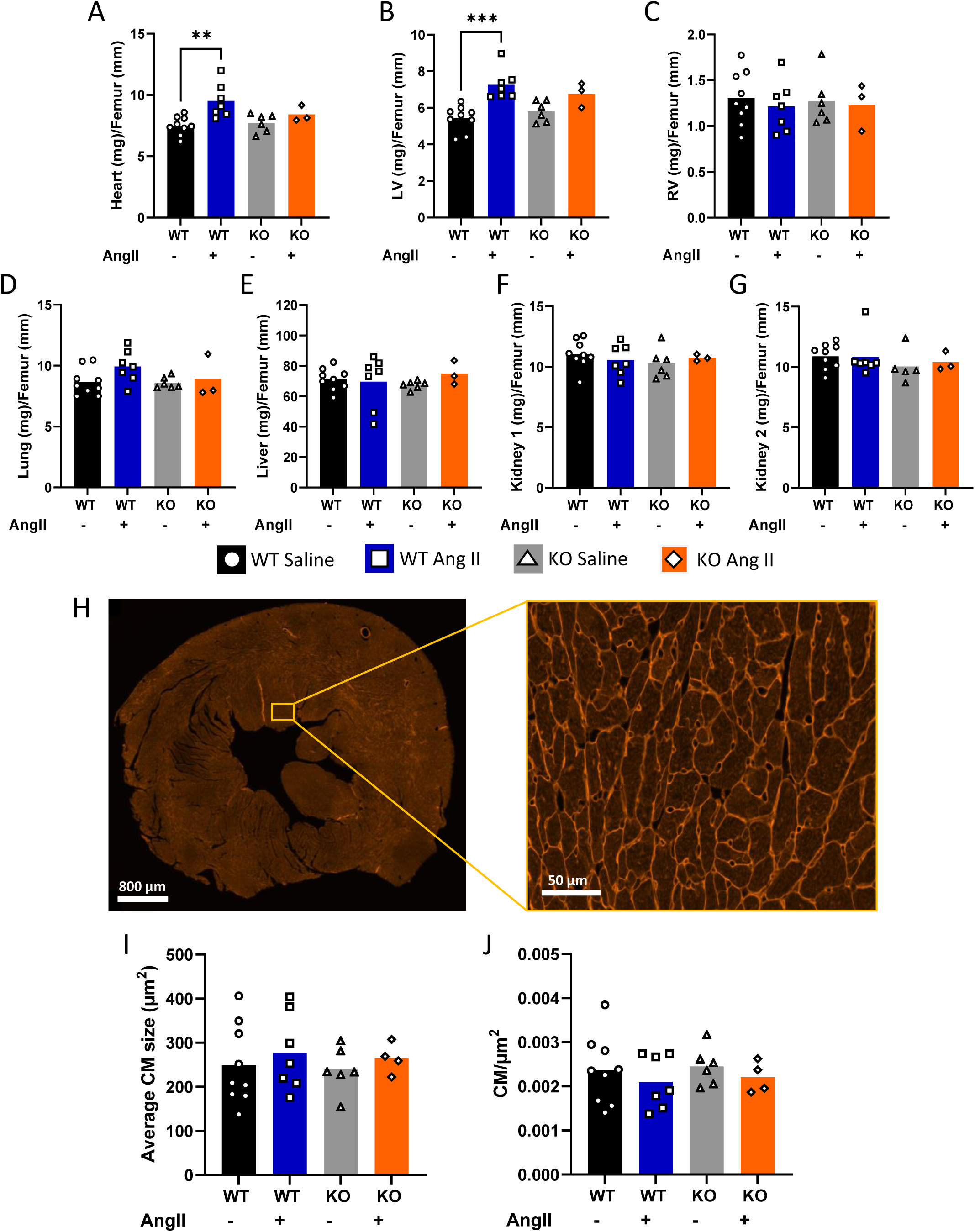
Comparison of organ weights and cardiomyocyte size in WT and *Mg23-*KO mice +/- AngII. Following treatment with either saline or AngII (1.1mg/kg/day), the **(A)** whole heart, **(B)** left ventricle (LV), **(C)** right ventricle (RV), **(D)** lung, **(E)** liver, **(F)** kidney 1 and **(G)** kidney 2 were weighed and normalised to femur length (mm). (WT saline 9 mice (black); WT AngII 7 mice (blue); KO saline 6 mice (grey); KO AngII 3 mice (orange)). **H)** Cell membrane in LV sections were stained with WGA and imaged at 20x. Representative image shown. **I)** Average cardiomyocyte (CM) size was measured using HALO. n ≥ 171 n ≤ 209 cells/slice. **J)** CM density was calculated as number of CMs/area (µm^2^). (WT saline 9 mice (black); WT AngII 7 mice (blue); KO saline 6 mice (grey); KO AngII 4 mice (orange)). Data is displayed as individual points with bar representing average. **p<0.01, ***p<0.001. 2-way ANOVA with post-hoc Tukey’s test.

### *Mg23*-KO hearts are protected against pressure-overload-induced fibrosis

Myocardial fibrosis is an important characteristic feature of pathological hypertrophy. Cardiac tissue slices prepared from the left ventricle of WT and *Mg23*-KO hearts were stained with Picrosirius Red to assess levels of fibrosis following saline or AngII treatment (Fig 3 A&B). The average fibrotic area in WT cardiac slices was significantly increased following AngII treatment compared to saline controls (10.54% ± 2.06% to 19.46% ± 1.84% (p <0.0001)). In contrast, no significant change in the level of fibrotic tissue in *Mg23*-KO hearts following AngII treatment compared to saline controls was observed (11.92% ± 1.77% control compared to 11.24% ± 1.4% AngII treated). Capillaries were evaluated using Isolectin-B4 stain, co-stained with Hoechst for scanning (Fig 3C). No significant difference was observed in vessel density when number of vessels were normalised to the total area analysed (Fig 3D). Blood vessels were subsequently separated into <30 µm^2^, 30 µm^2^ – 120 µm^2^ and >120 µm^2^ areas and expressed as a percentage of total blood vessel number. *Mg23*-KO hearts appeared to have a higher percentage of small vessels in saline control groups (<30 µm^2^) compared to WT hearts – 48.11% ± 2.69% in KO hearts and 39.68% ± 10.68% in WT (Fig 3E) however this difference was not significant (p = 0.076).

**Figure 3:**
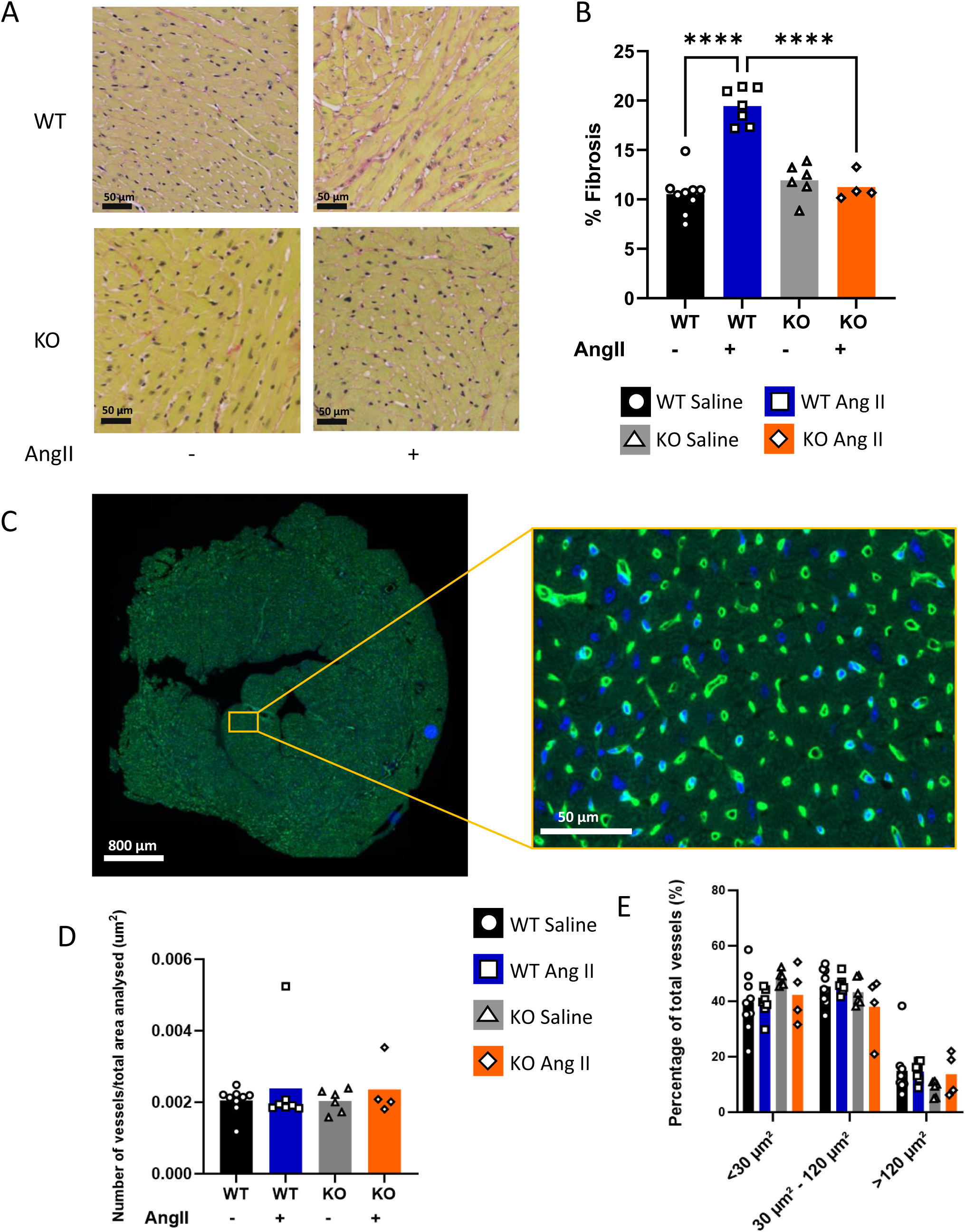
*Mg23-*KO mice are protected against AngII-induced fibrosis in left ventricle. **A)** Representative images of WT and *Mg23*-KO (KO) tissue slices. Picrosirius red staining was used to stain fibrillar collagen. **B)** % Fibrosis was measured for each slide. 10 squares were selected at random per slice and the average percentage fibrosis calculated. Data is displayed as individual points as average fibrosis per animal with bar representing group average. (WT saline 9 mice (black); WT AngII 7 mice (blue); KO saline 6 mice (grey); KO AngII 4 mice (orange)). **C)** Blood vessels were stained with isolectin (green) and Hoechst nuclear stain (blue). **D)** Number of blood vessels normalised to the total area analysed. **E)** Blood vessels were separated by size and percentage of each calculate. Data displayed as average percentage per animal with a bar representing the group average (WT saline 9 mice (black); WT AngII 7 mice (blue); KO saline 6 mice (grey); KO AngII 4 mice (orange)). ****p < 0.0001. 2-way ANOVA with post-hoc Tukey’s test.

### Dynamic cardiac functions following AngII treatment are preserved in *Mg23*-KO mice

We next examined the *in vivo* cardiac dynamics using pressure-volume catheters advanced into the left ventricle using a closed chest approach. Table 1 shows the cardiac functioning parameters and measurements for WT and *Mg23*-KO mice. Heart rate (HR) was consistent across all treatment groups (Fig 4A). In WT mice, AngII treatment induced an increase in the load-independent measurements of end-systolic pressure volume relationship (ESPVR, Fig 4I) and end-diastolic pressure volume relationship (EDPVR, Fig 4L) whereas in *Mg23-*KO mice, AngII did not induce an increase in ESPVR or EDPVR. Interestingly, significant differences were observed in the dV/dt max, a measure of rate of LV volume change, between the WT and *Mg23*-KO AngII treatment groups. Knock out of MG23 rescues hearts from decreased compliance seen by a reduction in ESPVR, EDPVR, V@dP/dt max (Fig 4J), dV/dt max (Fig 5A), dV/dt min (Fig 5B) and powMax/EDv (Fig 5C) in *Mg23*-KO animals compared to WT animals following AngII treatment.

**Figure 4:**
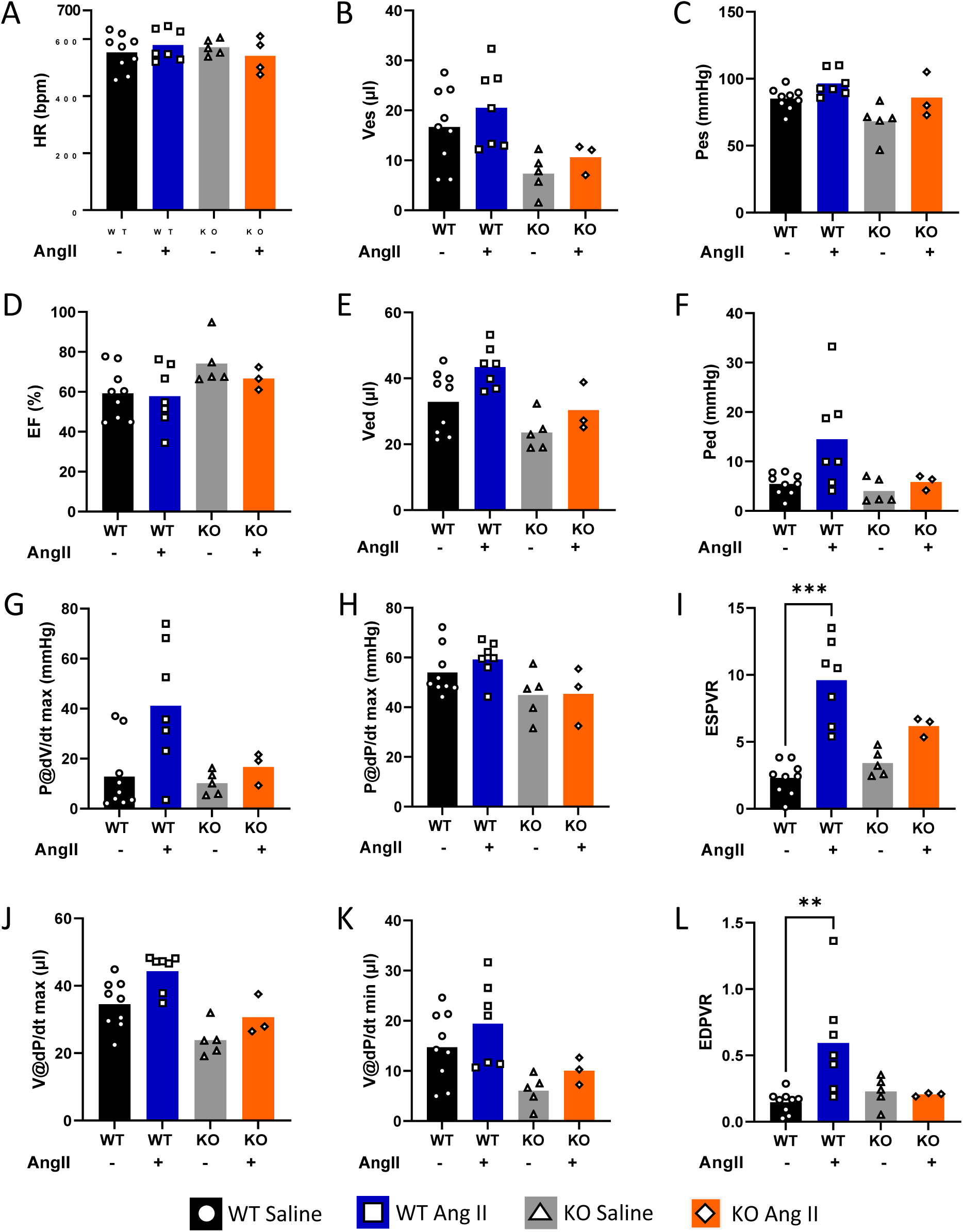
Cardiac function is preserved following AngII in *Mg23-*KO animals. **A)** Heart rate (HR; bmp), **(B)** Ves (µl), **(C)** Pes (mmHg), **(D)** ejection fraction (EF; %), **(E)** Ved (µl) and **(F)** Ped (mmHg) **(G)** P@dV/dt max (mmHg) **(H)** P@dP/dt max (mmHg), **(I)** ESPVR, **(J)** V@dP/dt max (µl), **(K)** V@dP/dt min (µl) and **(L)** EDPVR measured in WT saline (black; 9 mice), WT AngII (blue; 7 mice), *Mg23*-KO saline (grey; 5 mice) and *Mg23*-KO AngII (orange; 3 or 4 (HR) mice) animals. Data is displayed as an individual point per animal and a bar representing group average. **p<0.01, ***p<0.001. 2-way ANOVA with post-hoc Tukey’s test.

**Figure 5:**
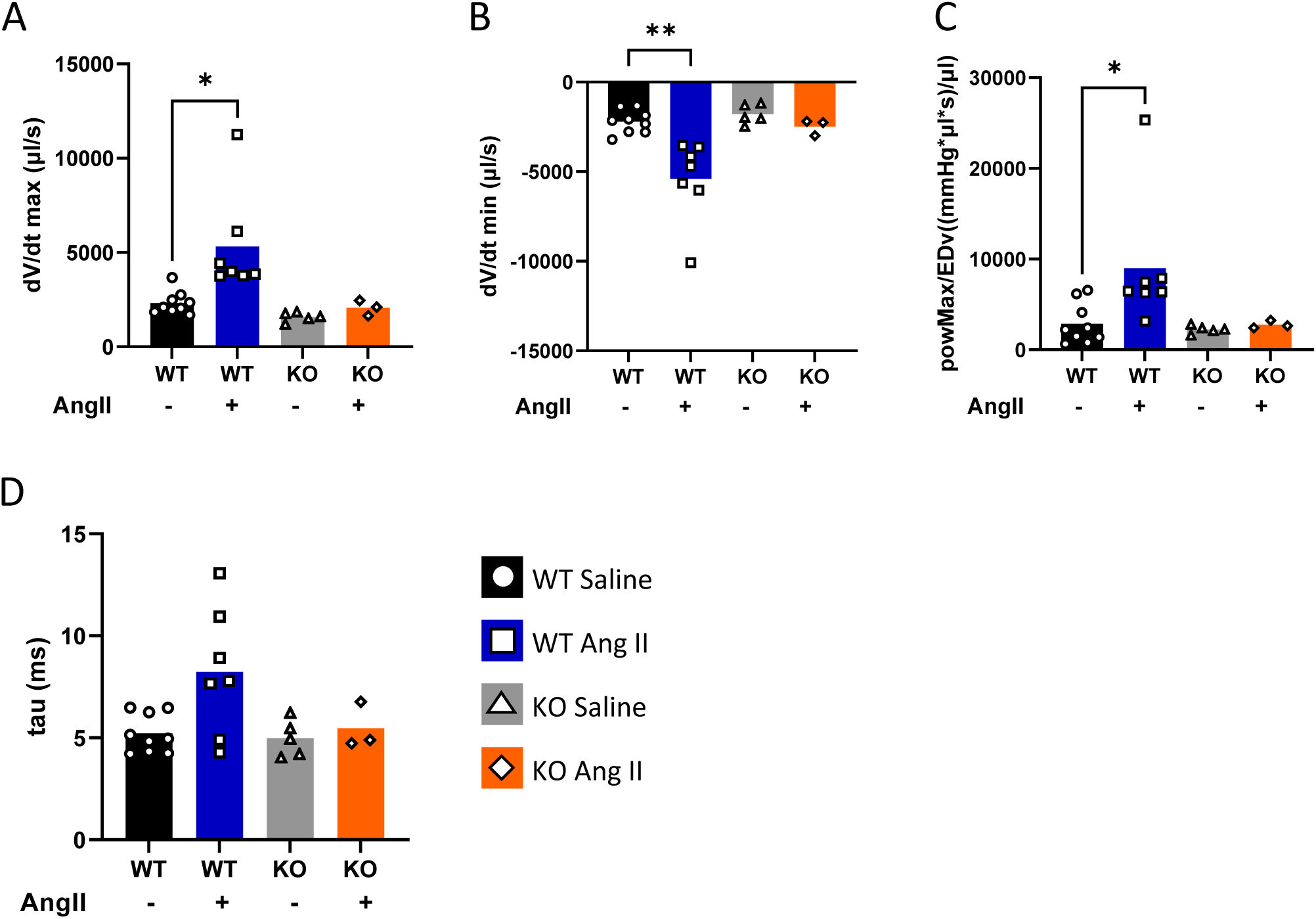
Cardiac compliance is preserved following AngII in *Mg23-*KO animals. **A)** dV/dt max (µl/s) **(B)** dV/dt min (µl/s) **(C)** powMax/EDV ((mmHg*µl*s)/µl) **(D)** tau (ms) measured in WT saline (black; 9 mice), WT AngII (blue; 7 mice), *Mg23*-KO saline (grey; 5 mice) and *Mg23*-KO AngII (orange; 3 mice) animals. Data is displayed as an individual point per animal and a bar representing group average. *p<0.05, **p<0.01. 2-way ANOVA with post-hoc Tukey’s test.

**Table 1:**
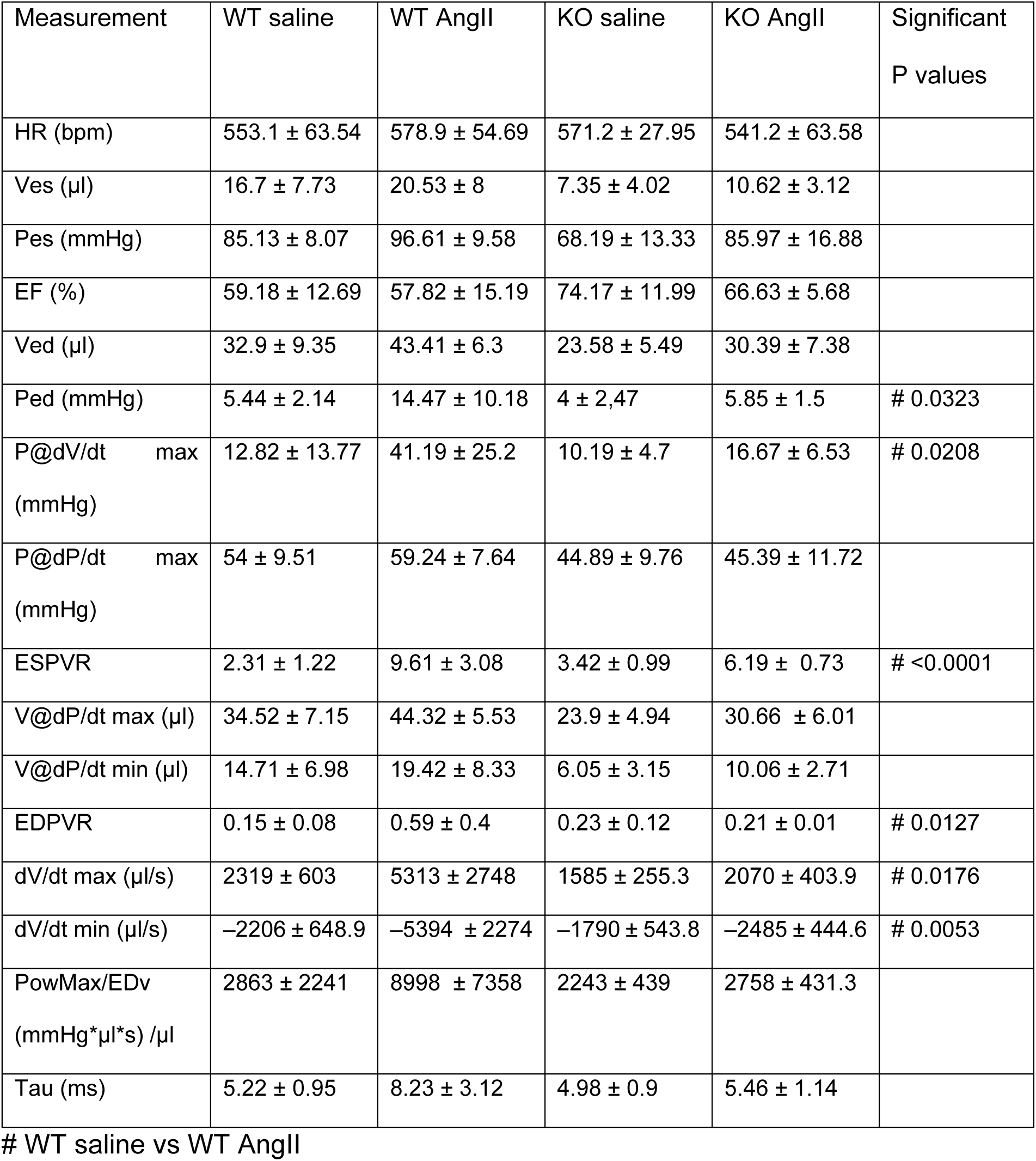
Cardiac Function parameters measured in WT and *Mg23-*KO mice treated with saline or Angiotensin II (AngII). Values shown are mean ± S.D.

| Measurement | WT saline | WT AngII | KO saline | KO AngII | Significant<br>P values |
| --- | --- | --- | --- | --- | --- |
| HR (bpm) | 553.1 ± 63.54 | 578.9 ± 54.69 | 571.2 ± 27.95 | 541.2 ± 63.58 | |
| Ves (μl) | 16.7 ± 7.73 | 20.53 ± 8 | 7.35 ± 4.02 | 10.62 ± 3.12 | |
| Pes (mmHg) | 85.13 ± 8.07 | 96.61 ± 9.58 | 68.19 ± 13.33 | 85.97 ± 16.88 | |
| EF (%) | 59.18 ± 12.69 | 57.82 ± 15.19 | 74.17 ± 11.99 | 66.63 ± 5.68 | |
| Ved (μl) | 32.9 ± 9.35 | 43.41 ± 6.3 | 23.58 ± 5.49 | 30.39 ± 7.38 | |
| Ped (mmHg) | 5.44 ± 2.14 | 14.47 ± 10.18 | 4 ± 2.47 | 5.85 ± 1.5 | # 0.0323 |
| P@dV/dt max<br>(mmHg) | 12.82 ± 13.77 | 41.19 ± 25.2 | 10.19 ± 4.7 | 16.67 ± 6.53 | # 0.0208 |
| P@dP/dt max<br>(mmHg) | 54 ± 9.51 | 59.24 ± 7.64 | 44.89 ± 9.76 | 45.39 ± 11.72 | |
| ESPVR | 2.31 ± 1.22 | 9.61 ± 3.08 | 3.42 ± 0.99 | 6.19 ± 0.73 | # <0.0001 |
| V@dP/dt max (μl) | 34.52 ± 7.15 | 44.32 ± 5.53 | 23.9 ± 4.94 | 30.66 ± 6.01 | |
| V@dP/dt min (μl) | 14.71 ± 6.98 | 19.42 ± 8.33 | 6.05 ± 3.15 | 10.06 ± 2.71 | |
| EDPVR | 0.15 ± 0.08 | 0.59 ± 0.4 | 0.23 ± 0.12 | 0.21 ± 0.01 | # 0.0127 |
| dV/dt max (μl/s) | 2319 ± 603 | 5313 ± 2748 | 1585 ± 255.3 | 2070 ± 403.9 | # 0.0176 |
| dV/dt min (μl/s) | -2206 ± 648.9 | -5394 ± 2274 | -1790 ± 543.8 | -2485 ± 444.6 | # 0.0053 |
| PowMax/EDv<br>(mmHg*μl*s) /μl | 2863 ± 2241 | 8998 ± 7358 | 2243 ± 439 | 2758 ± 431.3 | |
| Tau (ms) | 5.22 ± 0.95 | 8.23 ± 3.12 | 4.98 ± 0.9 | 5.46 ± 1.14 | |

### MG23 plays a key role in leakage of Ca^2+^ from sarcoplasmic reticulum Ca^2+^ stores

There is evidence to suggest that MG23 plays a role in the leakage of Ca^2+^ from sarcoplasmic reticulum stores^10,22^. Ca^2+^ store levels in fluo-4 loaded quiescent cardiomyocytes isolated from WT and *Mg23*-KO hearts were assessed by the application of caffeine. The peak *F/F_0_* response was significantly decreased in cardiomyocytes isolated from *Mg23*-KO animals from 5.79 ± 1.58 in WT cells to 4.06 ± 1.34 (p = 0.0002, fig 6D) showing a reduction in Ca^2+^ store levels in these cells. There was no significant change in the protein expression of RyR2, NCX1, SERCA2, Cav1.2, JP2 or CSQ2 in heart homogenates prepared from *Mg23*-KO animals compared to WT controls (Fig 6B). Although Ca^2+^ store levels were lower in *Mg23*-KO cells, significant increases in Ca^2+^ spark frequency (3.47 ± 1.42 sparks/ 100 µm/ sec to 7.97 ± 4.04 sparks/ 100 µm/ sec, (p = 0.0013, fig 6F), Ca^2+^ spark amplitude (1.66 ± 1.02 ΔF/F0 to 2.07 ± 2.31 ΔF/F0, p < 0.001, Fig 6G), time-to-peak (10.8 ± 15.34 ms to 15.99 ± 30.21 ms, p < 0.001, Fig 6H) and decay (Tau; 24.99 ± 50.08 ms to 37.18 ± 80.95 ms; p = 0.0017, Fig 6I) in *Mg23*-KO cardiomyocytes compared to WT controls were observed.

**Figure 6:**
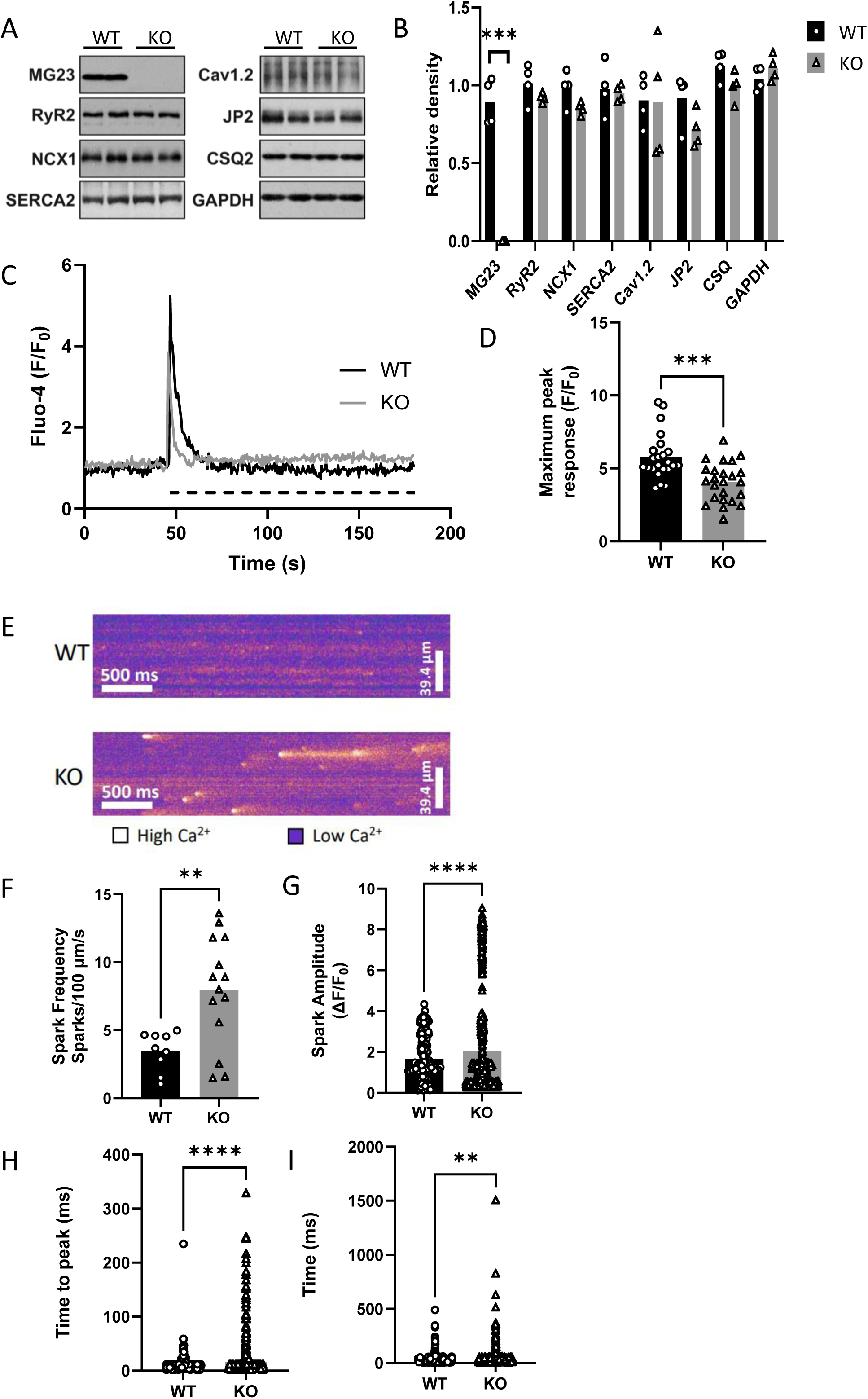
SR Ca^2+^ content and spark profiles are altered in *Mg23-*KO cardiomyocytes. **A)** Representative western blot images showing that expression of key Ca^2+^ -regulatory proteins is unaltered in *Mg23*-KO cardiac tissue. Lanes were loaded with 16 μg total protein. **B)** Bar graph showing quantification of data described in (A). Band densities were normalised to GAPDH loading controls. Data is displayed as individual cells with bar representing mean values (*n* = 4; 4 individual animals from each genotype). **C)** Representative trace showing Fluo-4 AM F/F_0_ of WT (black) and *Mg23*-KO (grey) isolated cardiomyocytes following addition of 10 mM caffeine. Caffeine addition illustrated on the graph as a dashed black line. **D)** Maximum peak response of Fluo-4 AM loaded wild-type (WT) and *Mg23*-KO (KO) isolated cardiomyocytes in response to 10 mM caffeine. Data is displayed as individual cells F/F_0_ with bar representing average peak response (n ≥ 20 cells from 2 animals). **E)** Representative line scan images showing spontaneous Ca^2+^ - spark events in cardiomyocytes isolated from WT (top) and *Mg*23-KO (bottom) mice. Graphs illustrating **(F)** Ca^2+^ - spark frequency, **(G)** spark amplitude, **(H)** time to peak and **(H)** decay (tau) measured from WT (black bars) and KO (grey bars) line scans. Individual points are shown with bars demonstrating average values (WT 289 sparks, 9 cells, 5 mice; *Mg23*-KO 1054 sparks, 11 cells, 5 mice). **p<0.01, ***p<0.001, ****p<0.0001. Multiple t-tests/unpaired t-test/unpaired t-test with Welch’s correction.

Nuclear Factor activated T cells transcription factors (NFATs) are linked to cardiovascular disease and hypertrophy (reviewed in^32^). Dephosphorylation of NFATs results in nuclear translocation where they influence gene transcription^33,34^. NFATc1, NFATc2 and NFATc3 are commonly linked to hypertrophy, heart failure and the Angiotensin II pathway^35–37^. We have developed a new methodology using multiplex immunofluorescence and AI analysis to assess cell specific NFAT regulation to study the emerging role of NFATs in cardiac pathology. Using left ventricular slices prepared from WT and *Mg23*-KO mice treated with AngII to induce pressure overload, we investigated the nuclear:cytosolic ratio of NFATc1, NFATc2 and NFATc3 by multiplex immunofluorescence. Cardiomyocyte positivity was confirmed by Troponin T staining, and cells were identified by cell membrane staining with WGA and nuclear Hoechst stain (Fig 7A). AI image analysis calculated the intensity of each target per cell in eight regions of interest. There was no significant difference in the ratio observed for NFATc1-3 (Fig 7). NFATc1 (Fig 7B, WT saline 1.29 ± 0.21; WT AngII 1.23 ± 0.15; KO saline 1.33 ± 0.22; KO AngII 1.39 ± 0.33), NFATc2 (Fig 7C, WT saline 1.92 ± 0.48; WT AngII 1.69 ± 0.37; KO saline 2.33 ± 0.68; KO AngII 1.83 ± 0.56) or NFATc3 (Fig 7D, WT saline 1.04 ± 0.08; WT AngII 1.02 ± 0.07; KO saline 1.04 ± 0.09; KO AngII 0.97 ± 0.04).

**Figure 7:**
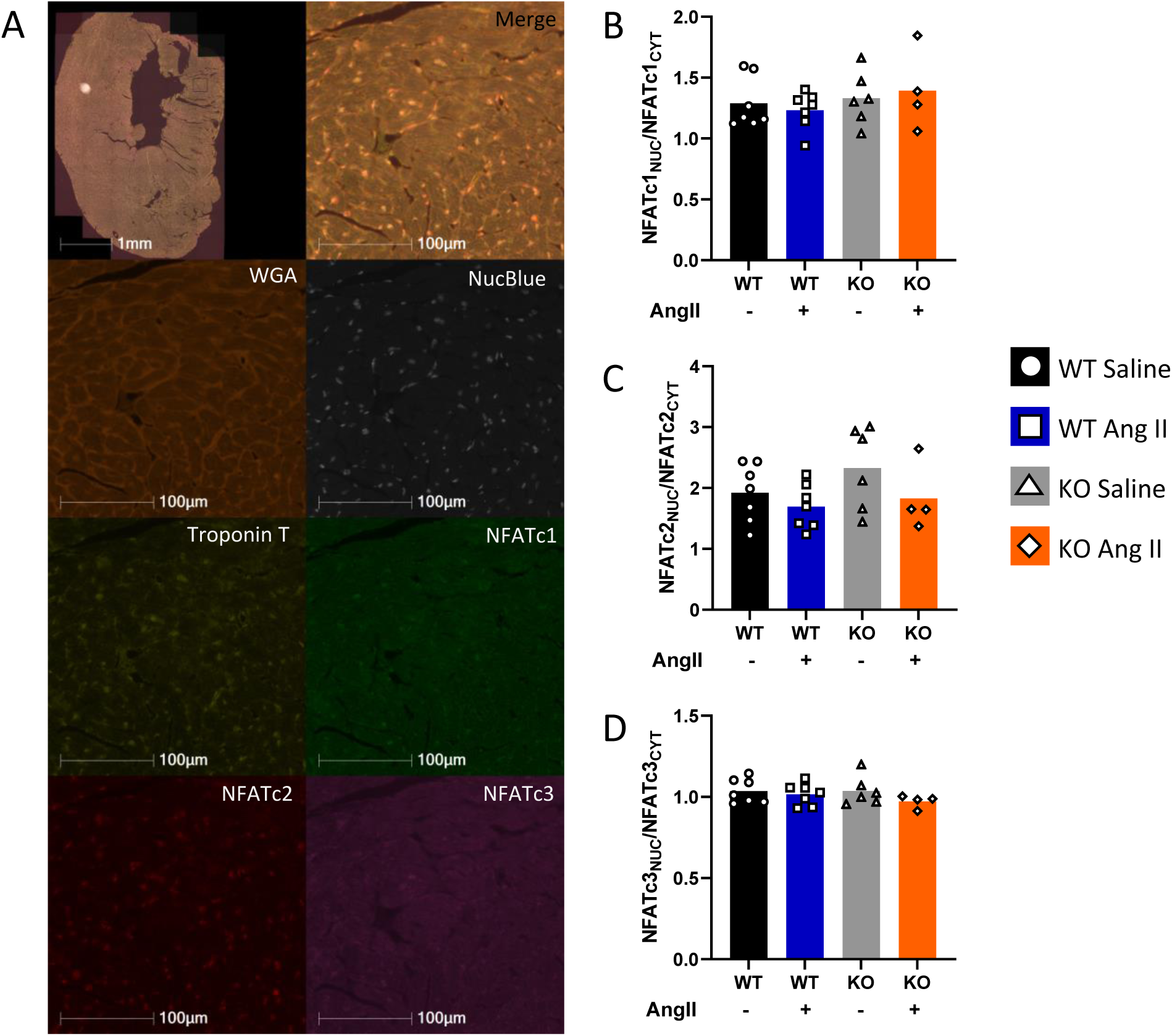
NFATs are not significantly altered in WT and *Mg23-*KO LV slices. **A)** Representative IF images of WGA (rhodamine), NucBlue (nuclear stain), Troponin T (cy3), NFATc1 (FITC), NFATc2 (cy5) and NFATc3 (Alexa750). Ratio of **(B)** NFATc1 (WT saline 1.29 ± 0.21; WT AngII 1.23 ± 0.15; KO saline 1.33 ± 0.22; KO AngII 1.39 ± 0.33) **(C)** NFATc2 (WT saline 1.92 ± 0.48; WT AngII 1.69 ± 0.37; KO saline 2.33 ± 0.68; KO AngII 1.83 ± 0.56) and **(D)** NFATc3 (WT saline 1.04 ± 0.08; WT AngII 1.02 ± 0.07; KO saline 1.04 ± 0.09; KO AngII 0.97 ± 0.04) in LV slices. 8 regions of interest were selected at random per slice and the intensity in nucleus and cytoplasm calculated using HALO. Data is displayed as individual points as average ratio per slice with bar representing group average. (WT saline 7 mice (black); WT AngII 7 mice (blue); KO saline 6 mice (grey); KO AngII 4 mice (orange)). 2-way ANOVA with post-hoc Tukey’s test.

## Discussion

In a cardiac pressure overload model, we have shown that knock out of MG23 protects the heart against left ventricular hypertrophy, altered compliance and cardiac fibrosis. We also provide the first direct evidence that MG23 plays a role in the leakage of Ca^2+^ from internal stores in cardiac tissue. These findings suggest that under pathological conditions MG23 contributes to cardiac dysfunction.

Ca^2+^ signalling plays a key role in the development of pathological cardiac hypertrophy^1^ but the exact nature of the subcellular Ca^2+^ pool that initiates hypertrophic signalling is not fully understood. Single channel studies have shown that MG23 is a Ca^2+^ permeable channel^21^ located to ER/SR and nuclear membranes^20^. The increased expression of MG23 in cardiac tissue that we observe following treatment with AngII may be highly relevant for hypertrophic transformation of the heart where functional perturbations in cellular Ca^2+^ homeostasis occur^38^. Hearts from *Mg23*-KO animals treated with AngII remained unchanged in size, whereas WT animals displayed increased heart weight due to increased left ventricular mass; a phenotype associated with significantly increased risk of heart failure and arrythmia^39,40^. The increase in heart mass seen in WT animals was not due to changes in cardiomyocyte size or cardiomyocyte density suggestive of pathological remodelling^41^. Indeed, our data show a significant increase in collagen fibre deposition in response to AngII treatment in WT animals that was not observed in *Mg23*-KO animals. We also observed a higher percentage of small vessels in cardiac tissue sections prepared from *Mg23*-KO hearts, suggestive of a higher rate of angiogenesis. Given that a lower microvascular density is associated with myocardial fibrosis in chronic cardiac diseases^42^ it is possible that the increase in microvascular density in *Mg23*-KO cardiac tissue protects against fibrosis in response to pressure overload through improved oxygen delivery in response to stress.

Pathological hypertrophy involves alterations in Ca^2+^ handling, increased oxidative stress, changes in excitation–contraction coupling, sarcomere dysfunction and metabolic and energetic remodelling^43–45^. These cellular and molecular changes can intersect to result in cardiomyocyte death and fibrosis. Fibrotic pathways are complex, encompassing cytokines, extracellular matrix proteins and matrix metalloproteinases (reviewed by^46^). While the specific fibrotic pathway linking MG23 to development of cardiac fibrosis is unknown, our data highlights MG23 as a potential therapeutic target for the prevention of cardiac fibrosis under cardiovascular stress. Interestingly, in HEK293 cells, MG23 has been implicated in apoptotic cell death^47,48^ and recent evidence has shown that MG23 may be part of the SND-pathway^49^. These data support a role for MG23 in ER stress responses suggesting that MG23 may be a driver in pathological remodelling responses. If MG23 plays a key role in modulating hypertrophic growth, this raises the prospect that therapies could be developed to target MG23 to modulate the hypertrophic response.

Measurements of ventricular performance demonstrated that pressure-overload induced changes in ventricular function in WT hearts. These changes were not observed in *Mg23*-KO hearts. This is consistent with *Mg23*-KO displaying limited changes in AngII induced cardiac remodelling and therefore being protected against hypertrophic transformational responses. In WT mice, measurements of contractility were increased following AngII treatment. These included the load-independent measurement of end systolic pressure response (ESPVR), which suggest that changing in cardiac function are independent of any vascular changes in blood pressure. Likewise, *Mg23*-KO mice were protected against the AngII-induced increase in diastolic pressure response (EDPVR). EDPVR represents passive filling properties of the left ventricle ^50^ which is increased after AngII treatment in line with enhanced myocardial stiffness often in relation with cardiac fibrosis^23^. In *Mg23*-KO hearts AngII was unable to increase passive tension. Interestingly, AngII increased other measurements of contractility (dV/dt max, PowerMax) in WT but not in *Mg23*-KO mice, evidencing hypercontractile state of the heart. At first the heart can adapt to these changes but over longer periods hypercontractile state will lead to heart failure^51,52^.

It has long been known that an undetectable leak of Ca^2+^ from the SR occurs during diastole (for review see^52^) and is thought to play an important role in sensitization of RyR2 for activation through Ca^2+^-induced Ca^2+^-release^7^ and cardiac protection against spontaneous Ca^2+^ release caused by SR Ca^2+^ overload^8^. Undetectable Ca^2+^ leakage plays an essential role in the maintenance of normal cardiac function but if this RyR2-independent leak becomes extensive it can become pathogenic leading to heart failure and the generation of fatal arrythmias^6,9,11,53^. The increased Ca^2+^ spark frequency and spark amplitude that we observe in isolated cells from *Mg23*-KO animals is suggestive of increased spontaneous RyR2 activity. RyR2 activity is regulated by SR luminal Ca^2+^ ^54–56^ suggesting that in *Mg23*-KO cells SR Ca^2+^ stores levels are elevated, but then rapidly deplete due to the increased activity of RyR2. This provides the first direct evidence that MG23 contributes to the leakage of Ca^2+^ from the SR. Although we cannot comment on the activity of SERCA in these cells, our data show that the expression of SERCA and other Ca^2+^ handling proteins is unaltered in cardiac tissue from *Mg23-*KO animals compared to WT animals. Our *in vivo* animal studies show that treatment with AngII results in increased expression of MG23 in cardiac tissue, linking MG23 to pathogenic responses.

MG23 is not only expressed on S/ER membranes it is also found in nuclear membranes, leading to the possibility that MG23 may influence gene expression through altered cellular Ca^2+^ dynamics. Activation of the calmodulin/Ca^2+^ dependent serine/threonine phosphatase calcineurin leads to dephosphorylation of NFATs resulting in their translocation to the nucleus where they induce alterations in gene expression (reviewed by^57^). Western blot analysis of NFAT phosphorylation status is notoriously variable^58^. Building on previous success utilising immunofluorescence to determine NFAT subcellular localisation in cell lines^59,60^, we developed a new methodology using multiplex immunofluorescence and AI analysis to assess the cardiomyocyte nuclear: cytosolic ratio of NFATs in ventricular slices. Although our data showed no difference in the nuclear:cytosolic ratio for NFATc1-3 between WT and *Mg23*-KO animals following treatment with AngII, it should be highlighted that there are multiple fibrotic pathways that drive cardiac hypertrophy including TGF-β, MAPK/p38 and YAP/TAZ and this warrants further investigation^61,62^ and reviewed by^1^.

## Conclusion

We propose that dysregulated Ca^2+^ leak driven through MG23 is a major player in cardiac fibrosis and resultant hypertrophic transformational responses.

## Nonstandard Abbreviations and Acronyms

AngII: Angiotensin II
BSA: Bovine serum albumin
Cav1.2: Calcium channel, voltage-dependent, L type, alpha 1C subunit
CM: Cardiomyocyte
CSQ2: Calsequestrin-2
EC coupling: Excitation-contraction coupling
EDPVR: End-diastolic pressure volume relationship
EF: Ejection fraction
ER: Endoplasmic reticulum
ESPVR: End-systolic pressure volume relationship
HR: Heart rate
IP3R: Inositol 1,4,5-trisphosphate (IP3) receptor
JP2: Junctophilin-2
LV: Left ventricle
MG23: Mitsugumin 23
NCX1: Sodium-calcium exchanger
NFAT: Nuclear Factor activated T cells
Ped: End-diastolic pressure
Pes: End-systolic pressure
PSR: Picro-Sirius Red
PV: Pressure volume
RV: Right ventricle
RyR2: Type-2 Ryanodine Receptor
SERCA: Sarco(endo)plasmic reticulum calcium ATPase
SR: Sarcoplasmic Reticulum
TSA: Tyramide signal amplification
Ved: End-diastolic volume
Ves: End-systolic volume
WGA: Wheat germ agglutinin

## Acknowledgments

We thank John O’Connor for assistance with preparation on cardiac slices for IF/IHC and help in carrying out the PSR staining. Travel grants awarded to A.M.D. from the Russell Trust (University of St Andrews), University of St Andrews School of Medicine, Biochemical Society and British Society for Cell Biology (Company of Biologists) allowed completion of key experiments.

## Author Contributions

C.S. & C.E.M. designed, performed and analysed the *in vivo* animal studies. I.H.U prepared the cardiac slices for IF/IHC and optimised the protocols under supervision of D.J.H. A.M.D & C.L.O performed and analysed the IF/IHC studies. A.M.D and G.B.R performed and analysed the *in vivo* cellular Ca^2+^ dynamic studies under the supervision of S.J.P. M.N, H.T and A.M.D. performed and analysed the biochemical experiments. S.J.P designed the project and wrote the manuscript with contributions from all authors.

## Supplemental Material

Supplemental Methods

Table S1

Figures S1 & S2

References ^23–25,63–65^

## Novelty and Significance

**What is known?**

- MG23 is a Ca^2+^ permeable ion channel located on sarco(endo)plasmic reticulum and nuclear membranes.
- Pathophysiological cellular Zn^2+^ levels, as observed in cardiac ischaemia, increase MG23 channel activity.
- MG23 modifies SR Ca^2+^ storage in skeletal muscle fibres.

**What new information does this article contribute?**

- MG23 contributes to SR Ca^2+^ leak in cardiomyocytes.
- In a murine model, *Mg23*-KO animals are protected against Angiotensin-II induced left ventricular hypertrophy/cardiac remodelling, including cardiac hypertrophy and left ventricular fibrosis.

